# *Variovorax paradoxus* alleviates bicarbonate-induced Fe limitation in cotton by altering the root-associated microbiome

**DOI:** 10.64898/2026.08.19.745819

**Authors:** Maruf Khan, Bishrant Pant, Ahmad H. Kabir

## Abstract

Alkaline and calcareous soils can induce iron (Fe) limitation in plants, yet the responses of root-associated microbial communities to beneficial rhizobacteria under these conditions remain poorly understood in cotton. Here, we investigated the effects of *Variovorax paradoxus* on plant performance, Fe nutrition, and root-associated microbiome dynamics in cotton exposed to bicarbonate-induced Fe limitation. In this study, *V. paradoxus* inoculation under bicarbonate-induced Fe limitation significantly improved photosynthetic parameters, growth parameters, and tissue Fe status. *V. paradoxus* inoculation partially reduced Fe-deficiency-induced root ferric-chelate reductase activity without a significant increase in net rhizosphere CAS-reactive Fe-chelating activity. This response may reflect a reduced requirement for maximal Strategy I activation following improved plant Fe status. Despite improved plant health, *V. paradoxus* reduced root C levels, suggesting altered belowground carbon utilization associated with bacterial inoculation and stress conditions. Split-root experiments showed greater improvements in chlorophyll score and shoot growth following bilateral than unilateral inoculation, although the effects of increased inoculum dose and root exposure could not be distinguished. In autoclaved soil, *V. paradoxus* did not improve plant growth under Fe deficiency, suggesting a possible role for resident soil microorganisms. Although bacterial alpha diversity remained unchanged, *V. paradoxus* significantly altered bacterial community composition and enriched *Cellvibrio* together with the fungal taxa *Funneliformis* and *Dominikia* under Fe limitation. Exploratory network analysis identified *Funneliformis* and *Serendipita* as highly connected fungal taxa associated with *V. paradoxus* treatment under Fe limitation, while core microbiome and indicator species analyses identified additional candidate microbial partners, including *Pseudomonas*, *Hydrogenophaga*, *Shinella*, and *Aquabispora*. Spearman correlation analysis further associated *Streptomyces* with root Fe accumulation and biomass, while *Epicoccum* and Sordariales were positively associated with CAS-reactive Fe-chelating activity in cotton exposed to bicarbonate-induced Fe limitation and inoculated with *V. paradoxus*. These findings highlight the potential of *V. paradoxus* to improve cotton Fe nutrition under bicarbonate-induced Fe limitation and identify root-associated microbial taxa whose functional contributions require further validation.

## 1. Introduction

Iron (Fe) is an essential micronutrient required for numerous physiological processes, including chlorophyll biosynthesis, photosynthetic electron transport, and respiration in plants (Kroh and Pilon, 2020). Although Fe is abundant in most soils, its bioavailability is severely restricted in calcareous and alkaline soils where ferric Fe (Fe^³⁺^) precipitates into insoluble forms, limiting plant uptake. Also, soil bicarbonate, poor aeration, and high concentrations of competing ions further exacerbate Fe deficiency by restricting Fe availability and impairing root Fe acquisition mechanisms (Kobayashi & Nishizawa, 2012; Briat et al., 2015). Consequently, Fe deficiency associated with calcareous and alkaline soils is one of the most widespread nutritional disorders affecting agricultural production worldwide, causing interveinal chlorosis, reduced photosynthesis, impaired growth, and substantial yield losses (Marschner, 2012; Kobayashi & Nishizawa, 2012; Briat et al., 2015). To cope with Fe limitation, dicot plants employ Strategy I responses. These include enhanced ferric-chelate reductase activity, increased proton extrusion to acidify the rhizosphere, activation of Fe² transporters, and changes in root architecture and exudation that improve Fe mobilization (Kobayashi & Nishizawa, 2012; Clemens and Weber, 2016). However, the efficiency of these adaptive responses varies substantially among plant species and genotypes. Consequently, plant genotypes vary widely in Fe-deficiency tolerance, with Fe-inefficient genotypes developing severe chlorosis and growth inhibition in calcareous soils (Zuo & Zhang, 2011).

Beneficial rhizosphere microorganisms have emerged as an environmentally friendly strategy to enhance plant nutrient acquisition and stress tolerance. Plant growth-promoting microorganisms enhance Fe availability through microbial siderophore, mineral solubilization, phytohormone synthesis, modification of root architecture, and changes in rhizosphere chemistry (Backer et al., 2018; Jacoby et al., 2017). Beyond their direct effects on plant physiology, plant-beneficial microorganisms can alter the indigenous root microbiome by selectively enriching taxa and modifying microbial associations that may support nutrient cycling and plant resilience under environmental stress. Among these beneficial microorganisms, *Variovorax paradoxus* is recognized for its remarkable metabolic versatility, efficient root colonization, and ability to interact with diverse members of the rhizosphere microbiome (Han et al., 2011). It is a non-spore-forming soil bacterium commonly found in the rhizosphere (Dodd et al., 2009). Previous studies have demonstrated that certain strains possess 1-aminocyclopropane-1-carboxylate (ACC) deaminase activity, produce indole-3-acetic acid (IAA), and colonize plant roots, thereby contributing to root development, nutrient acquisition, and tolerance to environmental stresses (Belimov et al., 2009; Chen et al., 2013). However, its role in improving Fe acquisition and regulating root microbiome assembly in plants under Fe-deficient conditions remains poorly understood.

Cotton (*Gossypium hirsutum* L.) is the world’s leading natural fiber crop and an important source of edible oil and livestock feed (Wu et al., 2022). Since cotton is extensively cultivated in calcareous and alkaline soils, indirect Fe deficiency frequently limits plant growth and fiber yield. Despite growing recognition of the role of root-associated microorganisms in plant nutrition, little is known about how *V. paradoxus* influences bacterial and fungal community composition in cotton experiencing bicarbonate-induced Fe limitation. Therefore, we investigated whether *V. paradoxus* improves cotton Fe nutrition under bicarbonate-induced Fe limitation and whether this response is accompanied by changes in the root-associated microbiome. We hypothesized that (i) *V. paradoxus* inoculation would improve cotton Fe nutrition, potentially without a corresponding increase in net rhizosphere CAS-reactive Fe-chelating activity, (ii) bilateral *V. paradoxus* inoculation would produce greater improvements in plant growth and chlorophyll status than unilateral inoculation, and (iii) *V. paradoxus* inoculation would be accompanied by changes in root-associated microbial community composition and the relative abundance of taxa potentially associated with stress recovery in cotton.

## 2. Materials and methods

### 2.1. Plant cultivation and growth conditions

Seeds of upland cotton (*Gossypium hirsutum* L.; USDA-GRIN accession SA-3928) were surface-sterilized with 2% sodium hypochlorite for 5 min, rinsed three times with sterile distilled water, and germinated in seedling trays at 25 °C for 48 h. Uniform seedlings were transplanted into pots containing 500 g of a soil mixture (field soil combined with commercial potting mix at a 1:2 ratio). Four treatment groups were established using NaHCO_3_-induced alkalization and *V. paradoxus* inoculation (1 × 10^8^ CFU/mL; ARS Culture Collection–USDA, B-1908) to study the effect of *V. paradoxus* on alkalinity-induced indirect Fe deficiency: (i) control (untreated soil, pH ∼6.5), (ii) - Fe (bicarbonate-treated soil, pH ∼7.8), (iii) -Fe+VP (bicarbonate-treated soil, pH ∼7.8, plus *V. paradoxus* inoculum), and (iv) +VP (untreated soil, pH ∼6.5, plus *V. paradoxus* inoculum). For plant inoculation, *V. paradoxus* was cultured in nutrient broth at 28 °C for 24 h on an orbital shaker (150 rpm). Following incubation, bacterial cells were harvested, washed, and resuspended in sterile distilled water to obtain a final concentration of 1 × 10 CFU mL ¹. For inoculation, 1 mL of the bacterial suspension was applied to the base of each seedling at the time of transplantation. Uninoculated control plants received an equivalent volume (1 mL) of sterile distilled water.

In addition, 50 mL of 15 mM NaHCO_3_ solution was applied weekly to bicarbonate-treated pots throughout the 6-week growth period to maintain alkaline conditions. Bicarbonate-induced Fe deficiency is widely used to simulate Fe chlorosis in alkaline and calcareous soils (Kabir & Bennetzen, 2024; Lucena et al., 2007). In this study, each treatment consisted of three independent biological replicates, with one plant per pot. The experiment was conducted twice independently to assess reproducibility. Data from one experiment were used for the figures and statistical analyses, while the second experiment confirmed the consistency of the observed treatment responses. An additional experiment was conducted using autoclaved soil under the same four treatment conditions (control, −Fe, −Fe+VP, and +VP) to evaluate the effects of *V. paradoxus* inoculation on cotton growth under sterilized soil conditions (Supplementary Fig. S1). Plants were grown for six weeks in a greenhouse under a completely randomized design with a 10-h light/14-h dark photoperiod, a photosynthetic photon flux density of approximately 250 μmol m ² s ¹, and a temperature of 25 ± 2 °C.

### 2.2. Molecular detection of inoculants in root samples

The presence of *V. paradoxus* in root samples was assessed using conventional PCR with species-specific primers. Root samples were gently rinsed twice with sterile phosphate-buffered saline (PBS), briefly vortexed to remove loosely attached soil particles, and washed twice with sterile distilled water. Genomic DNA was extracted from approximately 0.2 g of root tissue using the cetyltrimethylammonium bromide (CTAB) method (Clarke, 2009). DNA concentration and purity were determined using a NanoDrop ND-1000 spectrophotometer (Thermo Fisher Scientific, Wilmington, DE, USA), and DNA samples were normalized to equal concentrations before PCR. PCR amplification was performed using specific primers (forward: 5′-CAATCGTGGGGGATAACGC -3′; reverse: 5′-GGCCGCTCC ATTCGCGCA -3′). PCR products were separated by electrophoresis on a 1.5% agarose gel stained with GelRed and visualized under GelDoc™ EZ Imager (Bio-Rad Laboratories, Hercules, CA, USA).

### 2.3. Growth and physiological measurements

Chlorophyll content was determined non-destructively using a SPAD chlorophyll meter (SPAD-502 Plus, Konica Minolta, Japan). The photosynthetic performance index (Pi_ABS) was measured on the uppermost fully expanded leaves at three different positions using a portable FluorPen FP 110 (Photon Systems Instruments, Czech Republic). Before measurement, leaves were dark-adapted for 1 h. Furthermore, shoot height was measured from the stem base to the tip of the longest leaf using a ruler, whereas root length was measured from the crown to the tip of the longest root after gently washing the root system. After morphological measurements, shoots and roots were separated and immediately weighed to determine shoot fresh weight and root fresh weight using an analytical balance.

### 2.4 Elemental determination in plant tissues

Root samples were carefully excised, rinsed thoroughly under running tap water, immersed in 0.1 mM CaSO_4_ solution for 10 min to remove surface-bound ions, and subsequently washed with deionized water. Leaf samples were washed only with deionized water. Samples were placed in labeled paper envelopes and oven-dried at 75 °C for 72 h until a constant dry weight was achieved. Ground tissues were acid-digested, and Fe concentrations were determined using inductively coupled plasma optical emission spectrometry (ICP-OES; ICPE-9820, Shimadzu, Japan). Tissue C and N concentrations were measured using a FLASH Smart elemental analyzer (Thermo Fisher Scientific, USA) at the Center for Advances in Water and Air Quality, Lamar University.

### 2.5. Rhizosphere CAS-reactive Fe-chelating activity

Rhizosphere soil was collected by gently brushing soil adhering to the roots. Rhizosphere Fe-chelating activity was assessed using the Chrome Azurol S (CAS) assay (Alexander & Zuberer, 1991). Briefly, rhizosphere soil suspensions were prepared in sterile distilled water, and the supernatant was mixed with CAS reagent. After incubation at room temperature, absorbance was measured at 630 nm using a spectrophotometer. CAS-reactive Fe-chelating activity (%) was calculated as [(Ar − As)/Ar] × 100, where Ar and As represent the absorbance of the reference CAS reagent and sample, respectively.

### 2.6. Ferric chelate reductase activity in the roots

Fresh root tissue (approximately 0.1 g) was excised, thoroughly rinsed with deionized water, and briefly washed in 0.2 mM CaSO_4_ for 5 min. The roots were then transferred to 1 mL of assay solution containing 10 mM CaSO_4_, 5 mM MES buffer (pH 5.5), 0.1 mM Fe(III)-EDTA, and 0.3 mM Na_2_-BPDS (bathophenanthroline disulfonic acid disodium salt). Samples were incubated for 1 h at room temperature in the dark. A reagent blank containing assay solution without root tissue was included for each assay. Following incubation, the absorbance of the Fe² –BPDS complex was measured at 535 nm using a spectrophotometer. Ferric chelate reductase activity (FCR) was calculated using the molar extinction coefficient (ε = 22.14 mM ¹ cm ¹) and expressed as μmol Fe² g ¹ fresh weight min ¹ (Kabir et al., 2012).

### 2.7. Split-root experiments

To compare plant responses to unilateral and bilateral root exposure to *V. paradoxus*, cotton seedlings were grown for 2 weeks in autoclaved sterile soil to develop sufficient root systems for the split-root assay. The roots were gently divided into two comparable root portions and transplanted into paired compartments containing equal amounts of the same soil mixture. Six treatment combinations were established: SR1, control/control; SR2, −Fe/−Fe; SR3, −Fe/−Fe+VP; SR4, −Fe+VP/−Fe+VP; SR5, control/control+VP; and SR6, control+VP/control+VP. Each inoculated compartment received 1 mL of *V. paradoxus* suspension at 1 × 10 CFU mL ¹. Consequently, unilateral inoculation (SR3 and SR5) involved a total of 1 mL of bacterial suspension per plant, whereas bilateral inoculation (SR4 and SR6) involved 2 mL per plant, resulting in differences in both total inoculum dose and root exposure. Fe-deficiency treatments and irrigation were applied independently to each compartment, with precautions taken to prevent movement of inoculum or solution between compartments. Each treatment included three independent biological replicates. Plants were maintained under the greenhouse conditions described above and harvested after 4 weeks.

### 2.8. Amplicon sequencing and data processing

Root-associated microbial communities were characterized using Illumina amplicon sequencing targeting the bacterial 16S rRNA gene and fungal internal transcribed spacer (ITS) region. Root samples were washed to remove loosely attached soil particles but were not surface-sterilized before DNA extraction. Therefore, the sequenced microbial communities were classified as the root-associated fraction, potentially including both rhizoplane and endophytic microorganisms. Amplicon libraries were prepared by amplifying the bacterial 16S rRNA (V3V4) gene using the primer pair 338F (5′-ACTCCTACGGGAGGCAGCAG-3′) and 806R (5′-GGACTACHVGGGTWTCTAAT-3′), and the fungal internal transcribed spacer (ITS) region using the primer pair ITS1FI2 (5′-GAACCWGCGGARGGATCA-3′) and ITS2 (5′-GCTGCGTTCTTCATCGATGC-3′). Paired-end sequencing (2 × 250 bp) was performed on the Illumina NovaSeq 6000 platform.

Raw sequence reads were quality-filtered and adapter-trimmed using Cutadapt, followed by denoising, chimera removal, and inference of amplicon sequence variants (ASVs) using the DADA2 pipeline (Callahan et al. 2016). Reads assigned to chloroplast and mitochondrial DNA were excluded from the bacterial dataset before downstream analyses. Taxonomic assignment of bacterial ASVs (amplicon sequence variants) was performed against the SILVA 138 reference database (Quast et al. 2013), whereas fungal ASVs were classified using the UNITE database (Abarenkov et al. 2024). All downstream analyses were performed in R using the phyloseq (McMurdie & Holmes, 2013) and related packages. Alpha diversity was estimated using the observed ASVs and Shannon diversity indices after rarefaction to an even sequencing depth. Beta diversity was assessed using non-metric multidimensional scaling (NMDS) based on Bray–Curtis dissimilarity. Community differences between treatments were tested using permutational multivariate analysis of variance (PERMANOVA). The relative abundances of bacterial taxa were calculated after agglomerating ASVs at the phylum, class, order, family, and genus levels using the phyloseq package, and the top 15 most abundant taxa at each taxonomic rank were visualized as stacked bar plots based on relative abundance. Differences in taxon relative abundance among treatments were assessed using the Kruskal–Wallis test, and P-values were adjusted for multiple comparisons using the Benjamini–Hochberg false-discovery-rate.

### 2.9. Co-occurrence network analysis

Bacterial and fungal ASVs were separately agglomerated to the genus level using the phyloseq package, and genus abundances were transformed to relative abundance. Genera with a mean relative abundance <0.1% or zero variance across samples were excluded before analysis. A global microbial co-occurrence network was constructed using pairwise Spearman’s rank correlations among retained genera. P-values were adjusted for multiple testing using the Benjamini–Hochberg procedure, and correlations with |ρ| ≥ 0.60 and false-discovery-rate-adjusted P < 0.05 were retained to generate an undirected network in igraph, and isolated nodes were removed. Highly connected genera were ranked according to degree centrality in the global bacterial and fungal networks. Each genus was assigned to the treatment in which it exhibited the highest mean relative abundance, and the five highest-degree genera within each treatment-associated group were selected for visualization (Berry and Widder, 2014). Given the limited biological replication, the network analysis was considered exploratory, and network connectivity was not interpreted as evidence of direct microbial interactions, community stabilization, or functional contributions to Fe acquisition.

### 2.10. Core microbiome and indicator species analysis

Core microbiomes were identified separately for bacterial and fungal communities at the genus level. Core taxa were defined using ≥70% prevalence and ≥0.1% mean relative abundance to identify consistently occurring and sufficiently abundant genera while excluding rare taxa. These study-specific thresholds follow the established approach of combining prevalence and abundance criteria for core microbiome identification (Neu et al., 2021). Relative abundances of the core genera were calculated, and the twenty most abundant genera were compared among the treatment groups. Furthermore, indicator species analysis was performed at the genus level in R using the indicspecies package. Relative abundance data were used to identify microbial taxa significantly associated with each treatment group. Indicator values (IndVal) were calculated based on the specificity and fidelity of each genus to a given group, and statistical significance was assessed using 999 permutation tests. The top indicator taxa with significant (*P* < 0.05) indicator values were visualized as horizontal bar plots using ggplot2. Given the limited replication, indicator taxa were interpreted as candidate treatment-associated taxa requiring validation in larger independent datasets.

### 2.11. Correlation analysis between microbial genera and plant traits

Correlation analysis was performed in R using Spearman’s rank correlation to evaluate associations between microbial genera and plant growth traits. Genus-level relative-abundance data were used, and the 30 most abundant genera, based on mean relative abundance across all samples, were included in the analysis. Pairwise correlations were calculated between genus-level relative abundances and measured plant traits. P-values were adjusted using the Benjamini–Hochberg procedure, and associations with FDR-adjusted P < 0.05 were considered significant. Given the limited replication, these correlations were interpreted as exploratory. Correlation coefficients were visualized as clustered heatmaps using the pheatmap package in R, with hierarchical clustering applied to both genera and plant traits based on correlation similarity. Positive and negative correlations are represented by red and blue colors, respectively.

### 2.12. Statistical analysis

Data are presented as mean ± standard deviation (SD) of three independent biological replicates (*n* = 3) from one experiment. The independent repetition was used to confirm the reproducibility of the treatment responses and was not included in the statistical analyses. Growth and physiological experimental data were analyzed using R software (version 4.4.1). Differences among treatments were determined by one-way analysis of variance (ANOVA) with treatment as the fixed effect. When significant differences were detected (*P* < 0.05), treatment means were compared using Fisher’s least significant difference (LSD) test, and different lowercase letters indicate significant differences among treatments.

## 3. Results

### 3.1. Effects of *V. paradoxus* inoculation on plant growth

The aboveground phenotype and root systems showed visible differences among the four treatments (Fig. 1A). Plants grown under control and *V. paradoxus* inoculation exhibited larger shoots and more extensive root systems, whereas plants under -Fe showed reduced shoot growth and smaller root systems. Plants inoculated with *V. paradoxus* under Fe deficiency showed an intermediate phenotype, with greater shoot and root development than Fe-deficient plants but lower growth than control and +VP plants (Fig. 1A). PCR analysis detected *V. paradoxus* DNA in root samples from the −Fe+VP and +VP treatments, whereas no amplification was observed in the control and −Fe treatments (Fig. 1B). The SPAD chlorophyll index was significantly reduced under Fe deficiency compared with the Control (Fig. 1C). *V. paradoxus*-inoculated plants under Fe deficiency (-Fe+VP) significantly increased the SPAD value relative to the -Fe treatment but remained lower than the control. Plants inoculated with *V. paradoxus* under Fe-sufficient conditions exhibited SPAD values comparable to the control (Fig. 1C). Similarly, Pi_ABS decreased in the leaf under Fe deficiency compared with the control (Fig. 1D). However, *V. paradoxus* inoculation under Fe deficiency significantly increased Pi_ABS relative to -Fe, while the plants inoculated with *V. paradoxus* showed similar Pi_ABS to those of control plants (Fig. 1D). In this study, shoot height and shoot fresh weight declined significantly under Fe deficiency relative to the control (Fig. 1E-F). *V. paradoxus* inoculation significantly increased both parameters under Fe deficiency compared with the plants solely cultivated under Fe deficiency. Plants inoculated with *V. paradoxus* exhibited shoot growth comparable to the control (Fig. 1E-F). Further, root length and root fresh weight decreased under Fe deficiency relative to the control (Fig. 1G-H). Plants deprived of Fe but inoculated with *V. paradoxus* significantly improved both root traits compared to Fe-deficient plants. Plants solely inoculated with *V. paradoxus* showed similar root features to those of controls (Fig. 1G-H). In the autoclaved soil experiment, Fe deficiency significantly reduced SPAD values, shoot height, and shoot fresh weight compared with the control (Supplementary Fig. S1). However, *V. paradoxus* inoculation under Fe-deficient conditions did not significantly improve any of these parameters relative to uninoculated Fe-deficient plants. Under Fe-sufficient conditions, *V. paradoxus*-inoculated plants exhibited growth parameters comparable to those of the control (Supplementary Fig. S1).

**Fig. 1.**
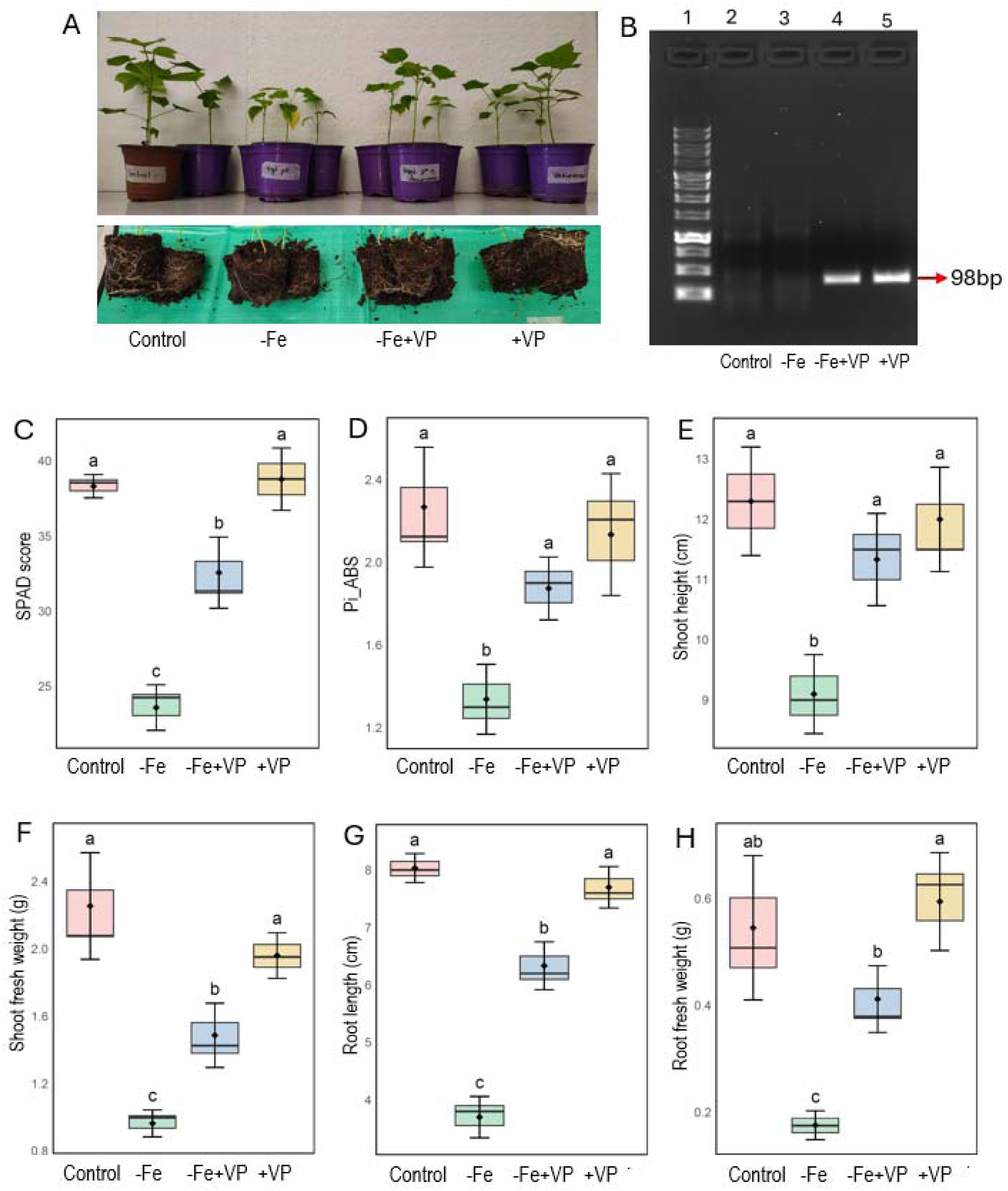
Effects of *V. paradoxus* (VP) inoculation on plant growth under Fe-deficient (-Fe) conditions. (A) Phenotype of whole plants (top) and root systems (bottom) grown under four treatments: control, Fe deficiency (-Fe), Fe deficiency with VP inoculation (-Fe+VP), and VP inoculation under Fe-sufficient conditions (+VP). (B) PCR-based detection of *V. paradoxus* DNA in root samples. Lane 1, 1 Kb DNA ladder; lane 2, control; lane 3, -Fe; lane 4, -Fe+VP; lane 5, +VP. (C–H) Effects of the treatments on (C) SPAD chlorophyll index, (D) photosynthetic performance index (Pi_ABS), (E) shoot height, (F) shoot fresh weight, (G) root length, and (H) root fresh weight. Data are presented as means ± SD from three biological replicates (*n* = 3). Different lowercase letters indicate significant differences among treatments according to one-way ANOVA followed by Fisher’s least significant difference (LSD) test (P < 0.05).

### 3.2. Changes in tissue elements and rhizosphere Fe-chelating activity

Bicarbonate treatment significantly reduced root and leaf Fe concentrations relative to the control (Fig. 2A). Inoculation with *V. paradoxus* under Fe deficiency (-Fe+VP) significantly increased Fe concentrations in both root and leaf relative to the -Fe treatment but remained lower than the control. The *V. paradoxus* treatment exhibited root Fe concentrations comparable to those of the control (Fig. 2A). N level did not differ significantly among treatments in both root and leaf of cotton plants (Fig. 2B). Interestingly, root C differed significantly among treatments. Root C concentration was significantly higher in both uninoculated treatments than in the corresponding plants inoculated with *V. paradoxus* (Fig. 2C). In leaves, C levels showed no significant differences among the treatment groups (Fig. 2C). In addition, rhizosphere CAS-reactive Fe-chelating activity differed significantly among treatments (Fig. 2D). Fe deficiency significantly increased Fe-chelating activity compared with the control. The −Fe+VP treatment did not differ significantly from the −Fe treatment in net rhizosphere CAS-reactive Fe-chelating activity, despite significantly higher root and leaf Fe concentrations. The VP+ treatment showed significantly lower CAS-reactive activity than both Fe-deficient treatments but was comparable to the control (Fig. 2D). Further, root ferric chelate reductase (FCR) activity increased significantly under Fe deficiency compared with the control (Fig. 2E). Inoculation with *V. paradoxus* significantly reduced FCR activity under Fe-deficient conditions compared to Fe-deficient plants, although activity remained higher than that of the control. Plants inoculated with *V. paradoxus* under Fe-sufficient conditions exhibited similar FCR activity to that of controls (Fig. 2E).

**Fig. 2.**
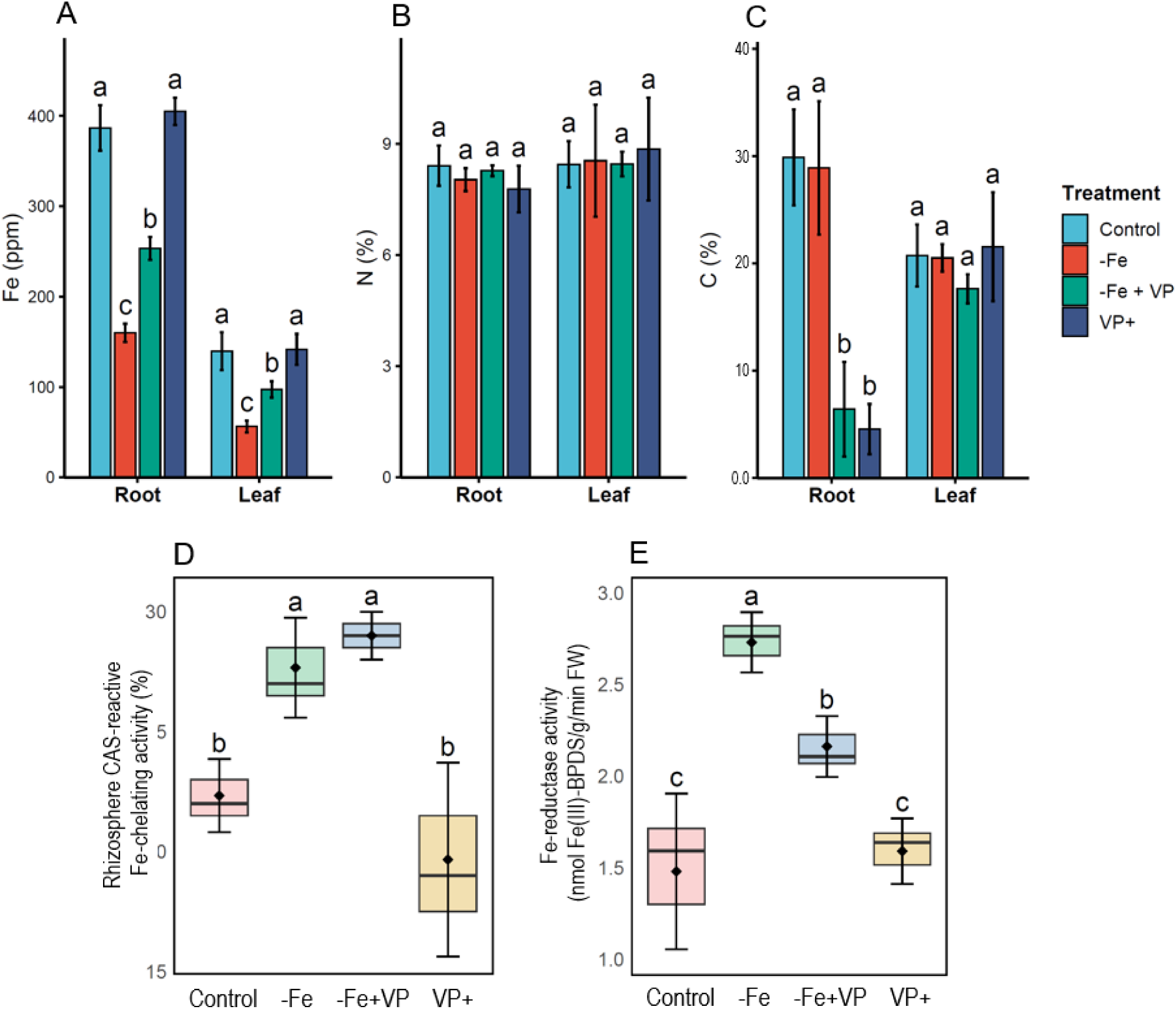
Effects of *V. paradoxus* (VP) inoculation on Fe acquisition under Fe-deficient (−Fe) conditions. Bar plots show (A) tissue Fe concentration, (B) tissue N content, and (C) tissue C content in roots and leaves, with error bars representing standard deviation (SD). Boxplots show (D) rhizosphere CAS-reactive Fe-chelating activity and (E) root ferric-chelate reductase activity, with boxes representing the interquartile range, horizontal lines indicating medians, and black diamonds indicating means. Data are presented as means ± SD from three biological replicates (n = 3). Different lowercase letters indicate significant differences among treatments according to one-way ANOVA followed by Fisher’s least significant difference (LSD) test (P < 0.05).

### 3.3. Effects of unilateral and bilateral split-root *V. paradoxus* inoculation on plant growth under bicarbonate-induced Fe limitation

The aboveground phenotype of split-root plants showed visible differences in plant growth (Fig. 3A). PCR analysis detected *V. paradoxus* DNA in inoculated root compartments, as indicated by the expected 98-bp amplicon, whereas no amplification was observed in uninoculated compartments (Fig. 3B). In split-root experiments, Fe deficiency imposed on both root compartments (SR2) significantly reduced SPAD values, shoot height, shoot fresh weight, and stem diameter relative to control plants having no stress in any of the compartments (Fig. 3C–F). When one root compartment was inoculated with *V. paradoxus* under Fe deficiency while the other remained uninoculated (SR3), all measured traits were significantly higher than those in SR2 but remained lower than those of the control. Bilateral VP inoculation under Fe deficiency (SR4) increased SPAD and shoot height relative to unilateral inoculation (SR3), restoring both traits to control-comparable levels, although shoot fresh weight remained lower than in the control (Fig. 3C, 3D and 3E). Under Fe-sufficient conditions, unilateral (SR5) and bilateral (SR6) inoculation served as controls to evaluate plant responses to V. paradoxus in the absence of Fe deficiency. Both treatments exhibited SPAD values, shoot height, and stem diameter comparable to the uninoculated control (SR1). Shoot fresh weight was also comparable between SR5 and SR1, whereas SR6 exhibited the highest shoot fresh weight among all treatments (Fig. 3C–F).

**Fig. 3.**
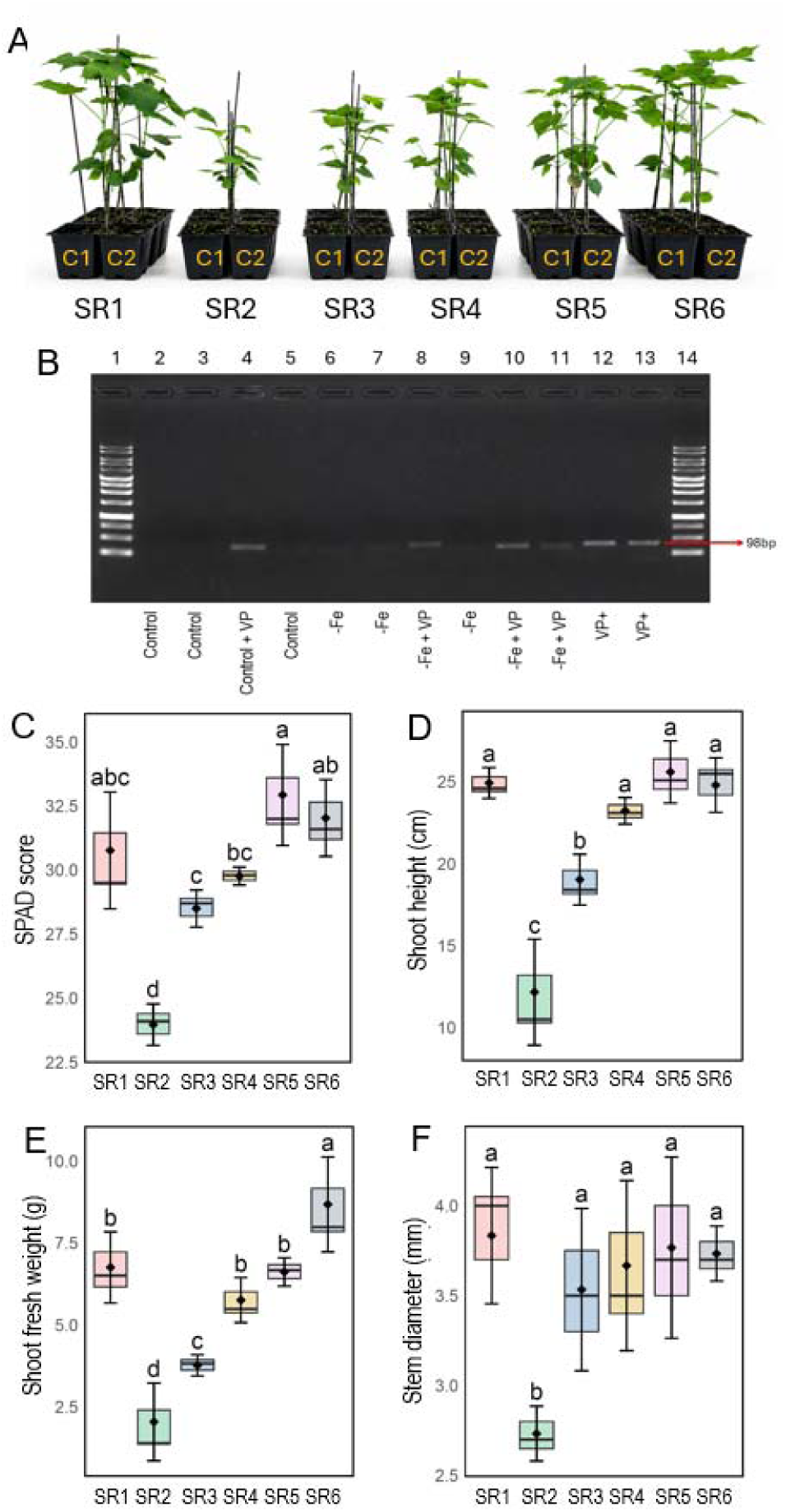
Growth performance of cotton plants having different split-root system cultivated in two compartments (C1: compartment 1, C2: compartment 2) in the presence or absence of *V paradoxus* (VP). SR1: control/control, SR2: -Fe/-Fe, SR3: -Fe/-Fe+VP, SR4: -Fe+VP/-Fe+VP SR5: control/control+VP, SR6: control+VP/control+VP) on aboveground phenotype (A), PCR-based detection of *V. paradoxus* DNA in root compartments (B), SPAD score (C), shoot height (D), shoot fresh weight (E), and stem diameter (F). Data are means ± SD from three biological replicates (*n* = 3), with letters indicating statistical differences based on one-way ANOVA followed by post hoc tests (*p* < 0.05).

### 3.4. Effects of *V. paradoxus* inoculation on root bacterial and fungal community dynamics

The root bacterial community composition differed among the four treatments, as shown by NMDS ordination based on Bray–Curtis dissimilarity (Fig. 4A). PERMANOVA indicated significant differences in bacterial community composition among treatments (R² = 0.472, *P* = 0.002), and the NMDS stress value was 0.075 (Fig. 4A). However, no significant differences were detected in bacterial alpha diversity among treatments based on either observed ASVs or the Shannon diversity index (Fig. 4B). The relative abundance of some dominant bacterial families varied among treatments (Fig. 4C). Community profiles were dominated by members of Comamonadaceae, Pseudomonadaceae, Rhizobiaceae, Sphingomonadaceae, and Xanthobacteraceae, with Methylococcaceae and Xanthobacteraceae differing significantly among treatments. Among these, Methylococcaceae and Xanthobacteraceae showed significant differential abundance among treatments (Fig. 4C). Similarly, the relative abundance of the dominant bacterial genera differed among treatments (Fig. 4D). Among the dominant genera, *Bradyrhizobium*, *Cellvibrio*, *Ideonella*, *Pseudoxanthomonas*, *Rhizobium*, and *Xylophilus* differed significantly among treatments, with *Cellvibrio* showing marked enrichment in −Fe+VP roots relative to −Fe roots (Fig. 4D). In particular, the abundance of *Cellvibrio* was substantially higher following *V. paradoxus* inoculation under Fe-limiting conditions than under Fe limitation alone. (Fig. 4D).

**Fig. 4.**
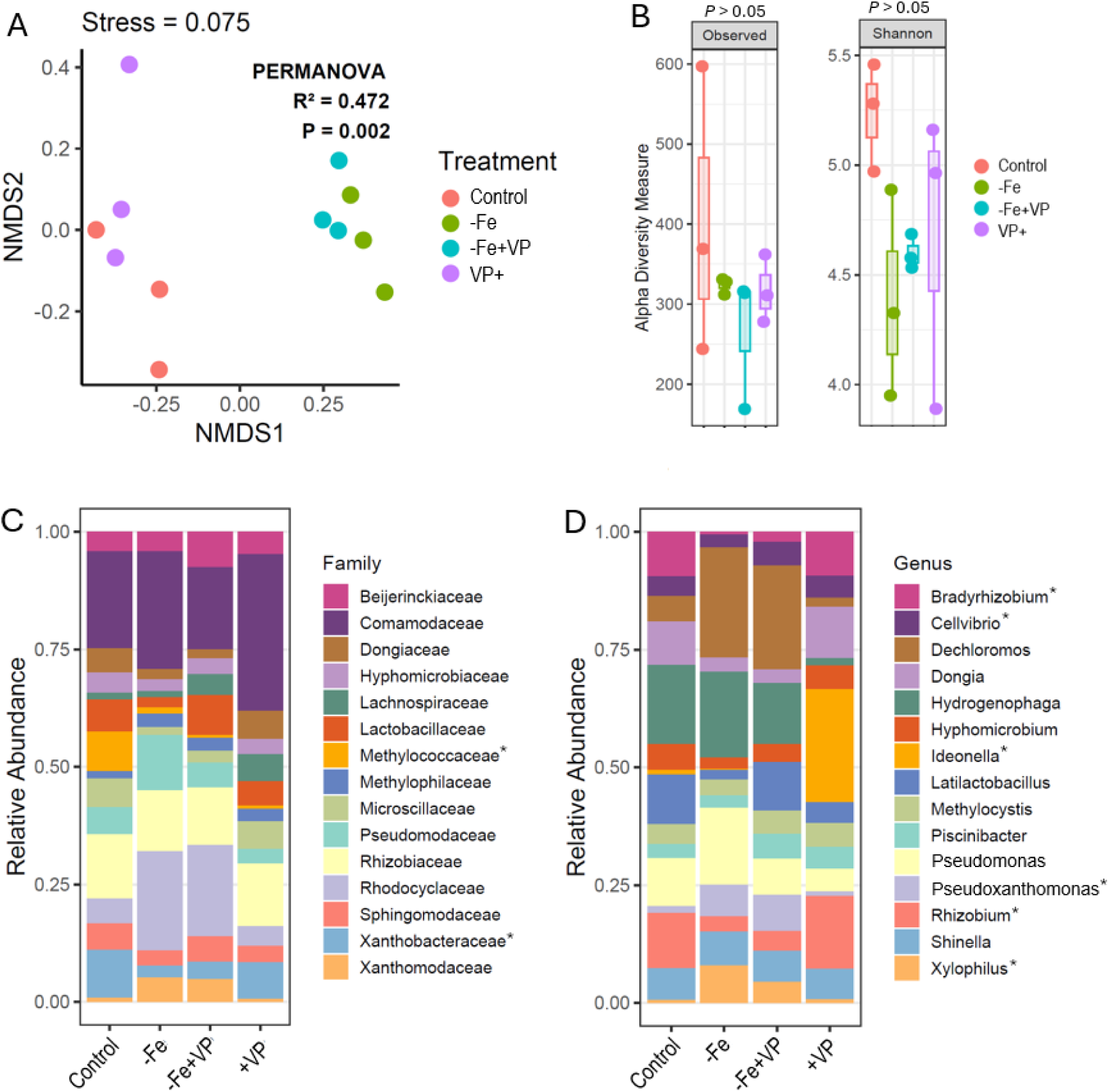
Root-associated bacterial community analysis under Fe deficiency and *V. paradoxus* (VP) inoculation. (A) Non-metric multidimensional scaling (NMDS) ordination based on Bray–Curtis dissimilarity showing bacterial community composition across the four treatments: control, Fe deficiency (-Fe), Fe deficiency with *V. paradoxus* inoculation (-Fe+VP), and *V. paradoxu* inoculation under Fe-sufficient conditions (VP+). PERMANOVA statistics (R² and *P* value) and NMDS stress value are indicated on the plot. (B) Alpha diversity of the bacterial community presented as observed ASVs (Observed) and Shannon diversity index for each treatment. Boxplots display the median, interquartile range, and minimum–maximum values, with individual points representing biological replicates. (C, D) Relative abundance of the dominant bacterial families and genera across treatments. Only the major families and genera are shown, and taxa marked with an asterisk (*) indicate statistically significant differential abundance.

In the case of fungi, NMDS ordination indicated no significant differences in root fungal community composition among treatments (PERMANOVA: R² = 0.332, P = 0.109; stress = 0.131) (Fig. 5A). Alpha diversity based on observed ASVs differed significantly among treatments, whereas the Shannon diversity index did not differ significantly (Fig. 5B). The relative abundance of the dominant fungal families varied among treatments (Fig. 5C). The major families identified across treatments included Acaulosporaceae, Aspergillaceae, Dictyosporiaceae, Didymellaceae, Diversisporaceae, Fuscosporaceae, Glomeraceae, Hydnodontaceae, Microascaceae, Nectriaceae, Orbiliaceae, Psathyrellaceae, Pseudteriellaceae, Rozellomycota, and Stachybotryaceae. Among these, Dictyosporiaceae, Glomeraceae, and Stachybotryaceae exhibited significant differential abundance among treatments. Similarly, the relative abundance of the dominant fungal genera differed among treatments (Fig. 5D). The predominant genera included *Acaulospora*, *Arthrobotrys*, *Boudiera*, *Dictyocheirospora*, *Diversispora*, *Dominikia*, *Funneliformis*, *Fusarium*, *Mucispora*, *Paramyrothecium*, *Phoma*, *Rhizophagus*, *Rozellomycota*, *Scedosporium*, and *Subulicystidium*. Among these, *Dictyocheirospora*, *Diversispora*, *Funneliformis*, *Paramyrothecium*, and *Subulicystidium* exhibited significant differential abundance among treatments (Fig. 5D). Also, the relative abundance of *Funneliformis* and *Dominikia* showed a significant enrichment in the roots of cotton plants inoculated with *V. paradoxus* under Fe deficiency compared to the plants solely cultivated with Fe deficiency (Fig. 5D).

**Fig. 5.**
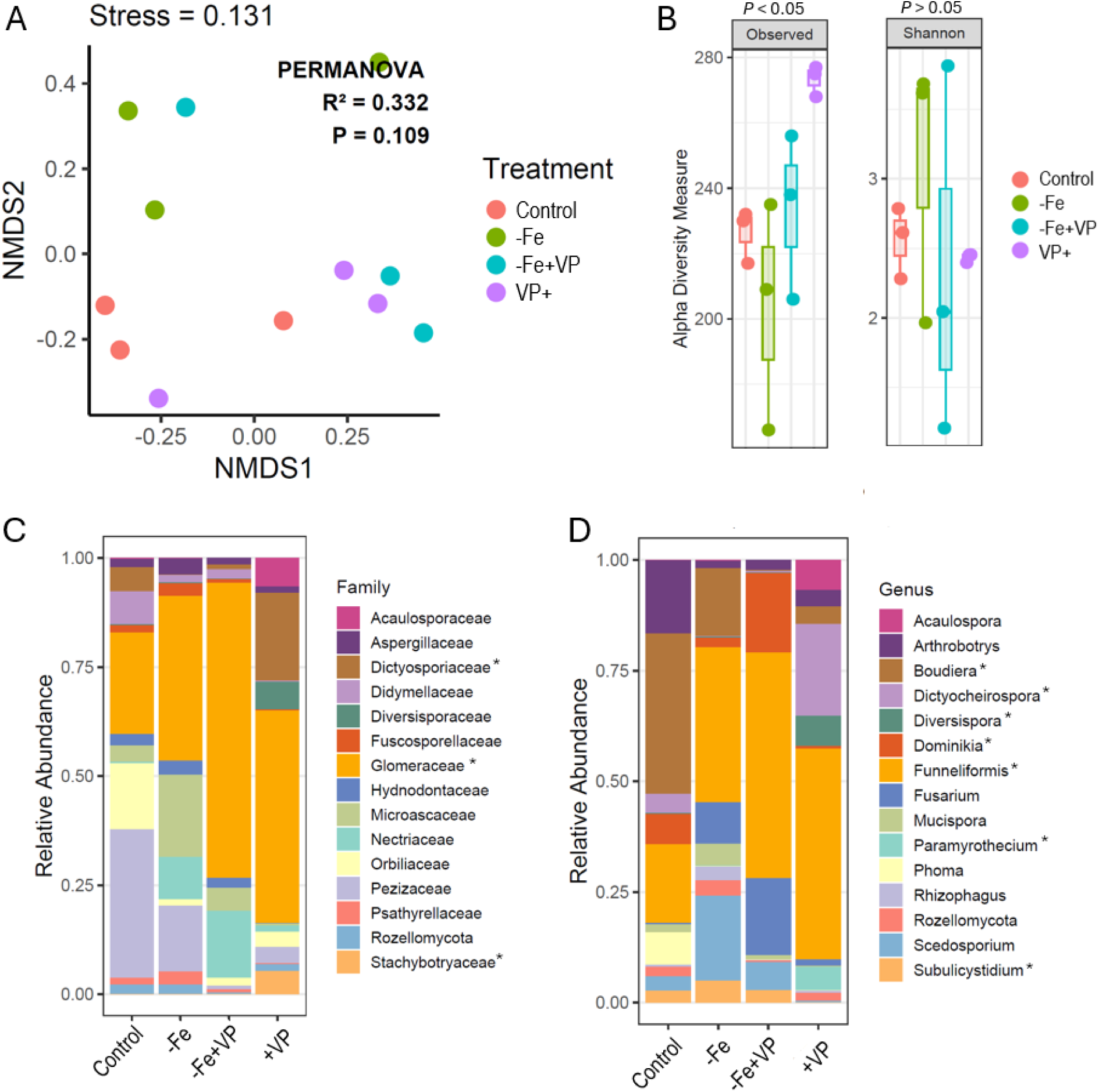
Root-associated fungal community analysis under Fe deficiency and *V. paradoxus* (VP) inoculation. (A) Non-metric multidimensional scaling (NMDS) ordination based on Bray–Curtis dissimilarity showing fungal community composition across the four treatments: control, Fe deficiency (-Fe), Fe deficiency with *V. paradoxus* inoculation (-Fe+VP), and *V. paradoxu* inoculation under Fe-sufficient conditions (VP+). PERMANOVA statistics (R² and *P* value) and NMDS stress value are indicated on the plot. (B) Alpha diversity of the fungal community presented as observed ASVs (Observed) and Shannon diversity index for each treatment. Boxplots display the median, interquartile range, and minimum–maximum values, with individual points representing biological replicates. (C, D) Relative abundance of the dominant fungal families and genera across treatments. Only the major families and genera are shown, and taxa marked with an asterisk (*) indicate statistically significant differential abundance.

### 3.5. Exploratory co-occurrence network identifies treatment-associated highly connected taxa

Exploratory global bacterial and fungal co-occurrence networks identified highly connected genera (candidate hub taxa) based on degree centrality (Fig. 6A–B). Each candidate hub taxon was assigned to the treatment in which it exhibited the highest mean relative abundance. Among bacteria, the top hub genera associated with the control group were *Asticcacaulis*, *Rhodomicrobium*, *Caulobacter*, *Phenylobacterium*, and *Hirschia* (Fig. 6A). Under Fe deficiency (-Fe), the dominant hub genera were *UBA6140*, *Azoarcus*, *C1-B045*, *Aquimos*, and *Pseudoxanthomonas*. The -Fe+VP treatment was characterized by *Chryseolinea*, *CL500-3*, *Arenibacter*, *SM1A02*, and *Sphingoaurantiacus*, whereas *Subgroup 10*, *Ohtaekwangia*, *Bradyrhizobium*, *Variovorax*, and *Ideonella* were identified as the major hub genera due to *V. paradoxus* inoculation. For fungal networks, the control treatment was associated with *Trichoderma*, *Cladophialophora*, *Penicillium*, *Cladosporium*, and *Boudiera* (Fig. 6B). Under Fe deficiency, the highly connected fungal genera were *Aspergillus*, *Paecilomyces*, *Chromelosporium*, *Alternaria*, and *Cercophora*. The -Fe+ *V. paradoxus* treatment was characterized by *Malassezia*, *Epicoccum*, *Cenococcum*, *Funneliformis*, and *Serendipita*, whereas *Xylomyces*, *Paramyrothecium*, *Acaulospora*, *Diversispora*, and *Atractiella* were identified as the principal hub genera in the *V. paradoxus+* treatment (Fig. 6B).

**Fig. 6.**
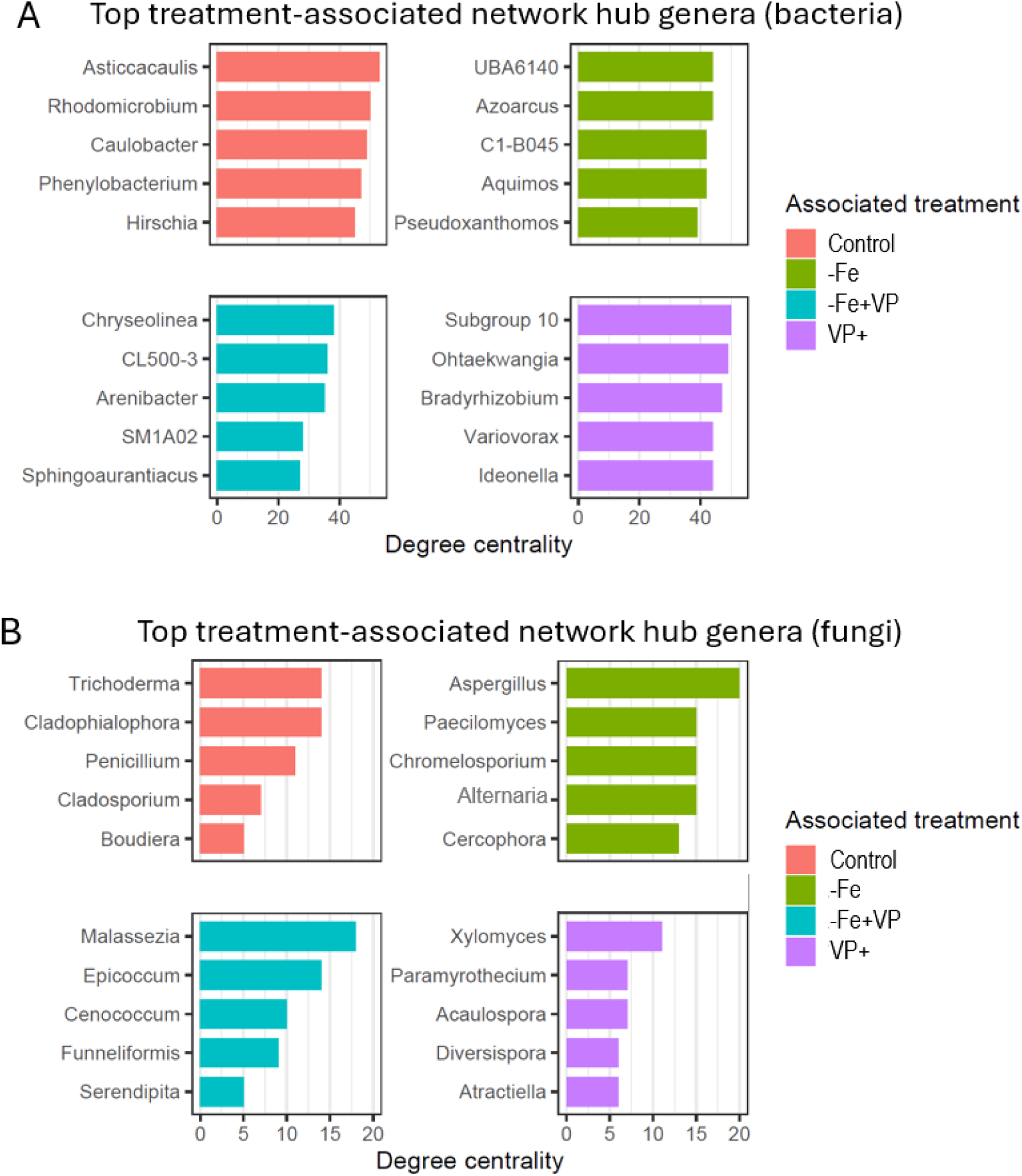
Treatment-associated highly connected genera identified from global bacterial and fungal co-occurrence networks. The five bacterial (A) and fungal (B) genera with the highest degree centrality are shown for each treatment association. Genera were assigned to the treatment in which they exhibited the highest mean relative abundance. Bar lengths represent degree centrality values calculated from the global bacterial and fungal co-occurrence networks. Color indicate these treatment associations rather than separate treatment-specific networks.

### 3.6. Core microbiome, indicator taxa, and plant trait associations

The relative abundance of the top core bacterial genera differed among treatments (Fig. 7A). The roots of control plants were dominated by *Dongia*, *Rhizobium*, *Hydrogenophaga*, Ohtaekwangia, *Bradyrhizobium*, and *Pseudomonas*. Under Fe shortage, the microbiome was dominated by *Dechloromonas*, *Hydrogenophaga*, *Pseudomonas,* and *Pseudoxanthomonas*. In the -Fe+ *V. paradoxus* conditions, the dominant core genera also included *Dechloromona*s, *Hydrogenophaga,* along with *Latilactobacillus*, *Pseudomonas,* and *Pseudoxanthomonas*. Plants solely inoculated with VP showed the dominance of *Dongia*, *Ideonella,* and *Rhizobium* (Fig. 7A). Indicator species analysis identified *Shinella*, *Ohtaekwangia*, *Dongia*, *Chryseolinea*, and *Cellvibrio* as taxa associated with the −Fe and/or −Fe+ *V. paradoxus* combination (Fig. 7B). In addition, *Bradyrhizobium*, *Rhizobium*, *Dongia*, and *Shinella* were identified as indicator taxa for the comparison between the control and *V. paradoxus*+ treatments (Fig. 7B).

**Fig. 7.**
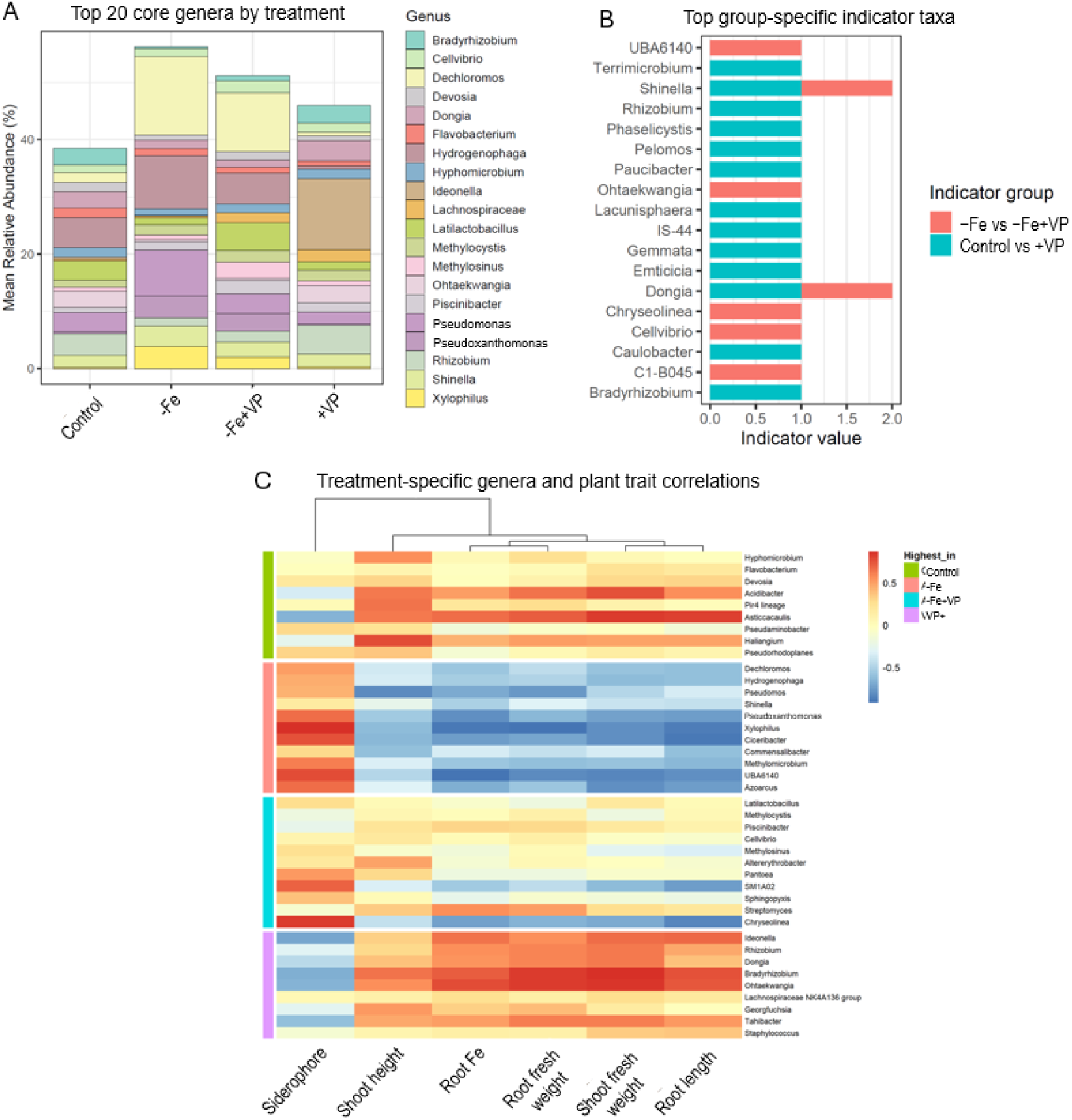
Core microbiome, indicator taxa, and plant trait associations of the root-associated bacterial community under different combinations of Fe deficiency and *V. paradoxus* (VP) inoculation. (A) Relative abundance of the top 20 core bacterial genera shared among the treatments. (B) Group-specific indicator bacterial genera identified by indicator species analysis. Bars represent indicator values for taxa associated with the treatment comparisons shown in the legend. (C) Heatmap showing Spearman’s correlation coefficients between treatment-specific bacterial genera and measured plant traits. Rows represent bacterial genera, columns represent plant traits, and colors indicate the strength and direction of the correlation. The colored annotation bar on the left indicates the treatment in which each genus was most abundant.

Spearman correlation analysis identified associations between microbial genera and plant traits. Cotton plants grown without Fe deficiency or *V. paradoxus* inoculation showed positive associations of *Acidibacter*, *Asticcacaulis*, and *Haliangium* with shoot height, root Fe concentration, root fresh weight, shoot fresh weight, and root length (Fig. 7C). Cotton plants grown under Fe deficiency showed strong positive associations of *Pseudoxanthomonas*, *Xylophilus*, *Ciceribacter*, and *Azoarcus* with siderophore production (Fig. 7C). Cotton plants grown under Fe deficiency with *V. paradoxus* inoculation showed strong positive associations of Streptomyces with root Fe concentration and root fresh weight, while *Pantoea* and *Chryseolinea* showed strong positive correlations with siderophore production. Cotton plants grown with *V. paradoxus* under Fe-sufficient conditions showed positive associations of *Rhizobium*, *Dongia*, *Bradyrhizobium*, and *Ohtaekwangia* with root Fe concentration, root fresh weight, shoot fresh weight, and root length (Fig. 7C).

In fungal analysis, cotton plants cultivated under control conditions were dominated by several core fungal genera, including *Arthrobotrys*, *Boudiera*, and *Funneliformis*, whereas Fe-deficient conditions were dominated by *Boudiera*, *Funneliformis*, and *Scedosporium* (Fig. 8A). Interestingly, Fe-deficient cotton plants inoculated with *V. paradoxus* were mostly dominated by *Dominikia*, *Funneliformis,* and *Fusarium*. However, *V. paradoxus* inoculation under Fe-sufficient conditions was dominated by *Dictyocheirospora* and *Funneliformis* (Fig. 8A). Indicator species analysis identified treatment-specific fungal taxa across pairwise comparisons (Fig. 8B). *Aquabispora* was the indicator taxon for −Fe vs −Fe+ *V. paradoxus*, whereas *Wettsteini* was the indicator taxon for −Fe vs −Fe+ *V. paradoxus* vs + *V. paradoxus*. The −Fe+ *V. paradoxus* treatment was characterized by *Mycothermus* and Lobulomycetales, while the comparison of −Fe+ *V. paradoxus* vs + *V. paradoxus* identified *Podospora*, Junewangiaceae, *Paecilomyces*, and *Pseudohypophila* as indicator taxa. The *V. paradoxus*+ treatment was distinguished by *Aspergillus*, *Paramyrothecium*, *Claroideoglomus*, and *Diversispora*, with *Diversispora* exhibiting the highest indicator value (Fig. 8B). Fungal taxa most abundant in the control treatment were positively associated with shoot fresh weight, root length, and root Fe concentration, including *Dictyosporium* and *Pluteus*. Genera enriched under Fe deficiency were primarily positively associated with siderophore production, including *Subulicystidium*, *Pseudallescheria* and *Lasiobolidium,* and *Chromelosporium* (Fig. 8C). Under Fe deficiency with *V. paradoxus* inoculation, *Funneliformis*, *Dominikia*, *Epicoccum,* and Sordariales showed strong positive associations with siderophore production. In contrast, fungal genera most abundant due to *V. paradoxus* inoculation, including *Dictyocheirospora*, *Diversispora*, *Acaulospora*, *Claroideoglomus*, and *Oehlia*, were positively associated with shoot fresh weight, root length, root Fe concentration, and root fresh weight (Fig. 8C).

**Fig. 8.**
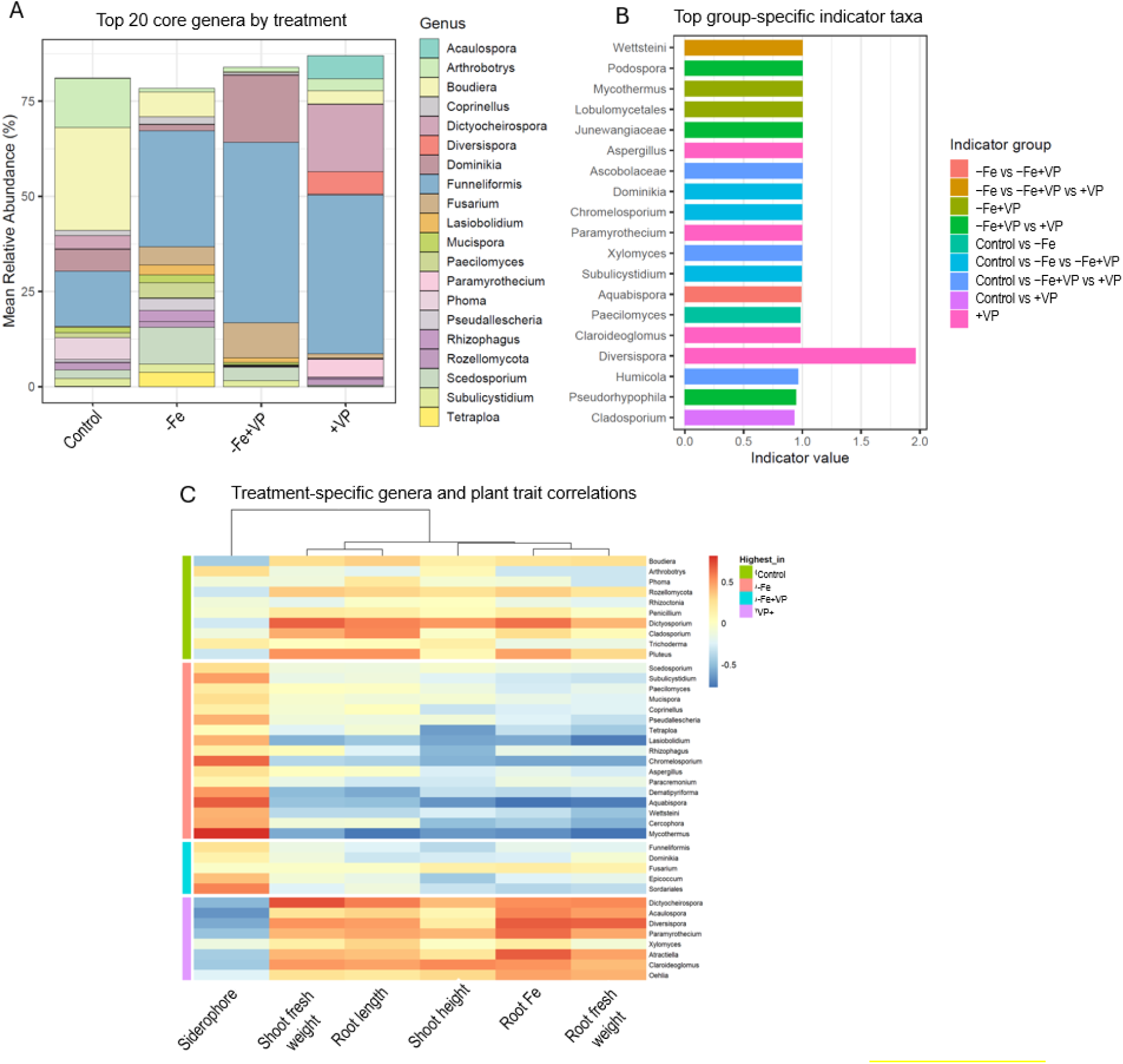
Core microbiome, indicator taxa, and plant trait associations of the root-associated fungal community under different combinations of Fe deficiency and *V. paradoxus* (VP) inoculation. (A) Relative abundance of the top 20 core fungal genera shared among the treatments. (B) Group-specific indicator fungal genera identified by indicator species analysis. Bars represent indicator values for taxa associated with the treatment comparisons shown in the legend. (C) Heatmap showing Spearman’s correlation coefficients between treatment-specific fungal genera and measured plant traits. Rows represent fungal genera, columns represent plant traits, and colors indicate the strength and direction of the correlation. The colored annotation bar on the left indicates the treatment in which each genus was most abundant.

## 4. Discussion

Beneficial rhizosphere microorganisms are increasingly recognized as sustainable tools to mitigate Fe deficiency symptoms and improve Fe acquisition and growth in plants (Thapa et al., 2025; Backer et al., 2018). Consistent with this concept, bicarbonate-induced alkaline conditions reduced cotton growth, photosynthetic performance, and tissue Fe accumulation, whereas inoculation with *V. paradoxus* substantially improved these traits and partially restored plant Fe status. This improvement occurred without an additional increase in CAS-reactive Fe-chelating activity and coincided with alterations in microbial community composition and the enrichment of selected bacterial and fungal taxa. Specifically, *V. paradoxus* enriched candidate microbial partners and was associated with distinct highly connected taxa while retaining a shared core microbiome. Together, these findings show that improved plant Fe status following *V. paradoxus* inoculation coincided with changes in bacterial community composition and enrichment of selected microbial taxa.

### 4.1. *V. paradoxus* improves photosynthetic performance and Fe nutrition under bicarbonate-induced Fe limitation

Inoculation with *V. paradoxus* improved root and leaf Fe accumulation, chlorophyll content, photosynthetic efficiency, and shoot and root growth under bicarbonate-induced alkaline conditions. Fe is indispensable for photosynthesis because it is required for chlorophyll biosynthesis, photosystem I, and enzymes involved in electron transport (Briat et al., 2015; Kalaji et al., 2017). The significant recovery of the photosynthetic performance due to *V. paradoxus* indicates restoration of functional photosynthetic machinery. Despite improved Fe status and photosynthetic performance, *V. paradoxus*-inoculated plants exhibited lower root C concentrations. However, the underlying physiological basis remains unclear because carbon allocation and turnover were not directly measured. Beneficial rhizobacteria increases the metabolic demand for recently fixed photoassimilates by stimulating root metabolism, nutrient acquisition processes, and microbial activity in the rhizosphere, thereby increasing carbon turnover in belowground tissues (Jones et al., 2009; Vives-Peris et al., 2020). Fe acquisition itself is an energy-intensive process that consumes carbohydrates derived from photosynthesis (Kobayashi & Nishizawa, 2012). Furthermore, the absence of significant differences in leaf carbon and nitrogen concentrations indicates that *V. paradoxus* maintained C and N homeostasis in the shoots. This response may reflect increased root C turnover or utilization associated with microbial colonization and Fe acquisition. However, direct measurements of rhizodeposition or carbon flux are required to confirm this interpretation.

Although *V. paradoxus* inoculation significantly improved tissue Fe concentrations under bicarbonate-induced Fe limitation, net rhizosphere CAS-reactive Fe-chelating activity did not differ significantly between inoculated and uninoculated Fe-deficient treatments. Importantly, this community-level assay cannot distinguish Fe-chelating compounds produced directly by *V. paradoxus* from those produced by resident microorganisms or other sources. Thus, unchanged net activity does not exclude siderophore production by *V. paradoxus*, which could be offset by changes in the activity of indigenous microorganisms. Improved Fe nutrition may involve direct bacterial effects, indirect changes in resident microbial activity, or plant-mediated responses. However, the relative contributions of these pathways remain unresolved and require pure-culture CAS assays and targeted functional experiments. Such possibilities are consistent with the metabolic versatility and rhizosphere interactions reported for *Variovorax* species (Ahmed & Holmström, 2014; Kramer et al., 2020; Han et al., 2020) but require direct experimental validation. Another possible explanation is that *V. paradoxus* enhanced the plant’s own Fe uptake capacity after Fe became mobilized in the rhizosphere. In Strategy I plants, Fe acquisition depends on rhizosphere acidification, ferric-chelate reduction, and Fe² transport (Marschner, 2012; Kobayashi & Nishizawa, 2012). Consistent with this well-established response, Fe-deficient cotton exhibited elevated root FCR activity. However, *V. paradoxus* inoculation partially reduced FCR activity while simultaneously increasing tissue Fe accumulation. This inverse relationship may reflect a reduced requirement for maximal activation of the plant’s Strategy I Fe-deficiency response following improved Fe status. Similar negative feedback regulation of Strategy I responses following improvements in plant Fe nutrition has been reported (Ivanov et al., 2012; Brumbarova et al., 2015). However, because rhizosphere Fe availability, Fe speciation, rhizosphere pH, and Fe transporter expression were not directly measured, this interpretation remains a plausible physiological explanation rather than a demonstrated mechanism. Together with the observed microbiome changes, these findings suggest that *V. paradoxus* may influence Fe acquisition through multiple physiological and rhizosphere-associated processes that require further investigation.

### 4.2. Bilateral *V. paradoxus* inoculation improves plant growth and chlorophyll status under bicarbonate-induced Fe limitation

The split-root experiment compared plant responses to unilateral and bilateral *V. paradoxus* inoculation under bicarbonate-induced Fe limitation. Bilateral inoculation produced greater improvements in SPAD and shoot growth than unilateral inoculation. However, because bilateral inoculation increased both the proportion of roots exposed and the total inoculum dose, the relative contributions of these factors could not be distinguished. The inoculation of *V. paradoxus* in only one root compartment partially improved plant traits; however, inoculation of both root compartments restored most physiological parameters compared with uninoculated Fe-deficient plants. Similar localized effects have been reported in plants for nutrient-solubilizing microorganisms (Kabir et al., 2024; Santoyo et al., 2021). The partial recovery following unilateral inoculation indicates that inoculation of a portion of the root system can influence whole-plant performance, although the present experiment does not distinguish local microbial effects from potential systemic responses. Plant responses to nutrient deficiency generally involve both local sensing at the root surface and long-distance shoot-to-root signaling that coordinates nutrient acquisition throughout the plant (Krouk et al., 2011; Giehl et al., 2014).

Further, *V. paradoxus* inoculation significantly increased both root length and root fresh weight compared with uninoculated Fe-deficient plants, indicating that improved Fe nutrition was accompanied by enhanced root system development. Localized microbial activity may also explain the pronounced improvement in root growth observed in fully inoculated plants. Rhizobacteria can alter root-zone pH, stimulate secretion of organic acids, and modify root exudation patterns, thereby creating favorable microenvironments for nutrient acquisition (Philippot et al., 2013; Jacoby et al., 2017). Such processes commonly occur near the root surface and may therefore depend on the extent of root–microbe contact. Although the main experiment revealed changes in root-associated microbial community composition following *V. paradoxus* inoculation, microbiome analyses were not performed on split-root samples. Therefore, the greater growth response following bilateral inoculation cannot be directly attributed to microbiome changes. Furthermore, tissue Fe concentrations and ferric-chelate reductase activity were not measured in the split-root experiment, limiting conclusions about the underlying Fe-acquisition mechanisms. These observations indicate that bilateral inoculation produced greater recovery than unilateral inoculation. However, because bilateral inoculation also increased the total inoculum dose and proportion of roots exposed to *V. paradoxus*, the contributions of these two factors cannot be separated.

### 4.3. *V. paradoxus* alters root microbial community composition under bicarbonate-induced Fe limitation

Plant growth-promoting microorganisms function not only through their own metabolic activities but also by reshaping the indigenous root microbiome into communities that better support plant nutrition and stress adaptation (Compant et al., 2019; Song et al., 2026). Although bacterial alpha diversity and fungal Shannon diversity remained unchanged, fungal observed richness differed among treatments, and bacterial community composition was significantly altered in cotton. These observations indicate that the beneficial effects of *V. paradoxus* coincided with changes in bacterial community composition rather than with increased bacterial diversity. This pattern is increasingly recognized in plant microbiome studies, where beneficial microorganisms influence the identity and relative abundance of community members rather than simply increasing microbial diversity (Toju et al., 2018; Taheri et al., 2025). Thus, improved Fe nutrition was more closely associated with changes in microbial composition and taxon abundance than with a general increase in microbial diversity.

*Cellvibrio* and *Pseudoxanthomonas* were among the bacterial genera that differed across treatments, with *Cellvibrio* showing pronounced enrichment following *V. paradoxus* inoculation under Fe deficiency. These taxa represent candidate community members potentially associated with nutrient turnover, although their functional contribution in the present system remains unknown. Species of *Cellvibrio* are recognized for producing diverse carbohydrate-active enzymes capable of degrading plant-derived polysaccharides and organic matter (Zhang et al., 2020; Gardner, 2024). Enhanced decomposition of organic substrates may increase the availability of low-molecular-weight organic compounds capable of chelating Fe or stimulating microbial metabolism in the rhizosphere. Similarly, *Pseudoxanthomonas* species participate in nutrient cycling and degradation of complex organic compounds (Peng et al., 2024). In contrast, fungal richness varied among treatments, whereas overall fungal community composition remained relatively stable, suggesting a weaker community-level response than that observed for bacteria. Fungal communities generally exhibit slower turnover rates and stronger dependence on long-term soil characteristics than bacterial communities (Bahram et al., 2018). Nevertheless, *V. paradoxus* significantly enriched the abundance of arbuscular mycorrhizal fungi (AMF) such as *Funneliformis* and *Dominikia* under Fe deficiency with *V. paradoxus*. Both genera belong to Glomeromycotina, whose members have been reported to facilitate plant nutrient acquisition through extraradical hyphal networks (Umer et al. 2025). Although AMF are traditionally associated with phosphorus uptake, increasing evidence demonstrates that they also facilitate micronutrient acquisition, including Fe, Zn, and Mn, particularly in calcareous soils where nutrient mobility is restricted (Lehmann & Rillig, 2015). The increased relative abundance of *Funneliformis* and *Dominikia* following V. paradoxus inoculation identifies these genera as candidates for further investigation. However, their contributions to Fe acquisition and potential interactions with *V. paradoxus* were not directly evaluated. Similar bacterial–fungal cooperation has recently been recognized as a fundamental mechanism governing microbiome-mediated plant resilience under nutrient stress (Lee et al., 2022). Therefore, their enrichment should be interpreted as a treatment-associated change rather than evidence of functional cooperation.

The differences among taxa identified by microbiome analyses may reflect variation in ecological niches, treatment responses, and analytical criteria. Enrichment of *Cellvibrio* may indicate changes in organic matter turnover, whereas the arbuscular mycorrhizal fungi *Funneliformis* and *Dominikia* may contribute to nutrient acquisition (Shi et al., 2021). The persistence of *Pseudomonas* and *Hydrogenophaga* suggests their roles as prevalent community members, while *Shinella* and *Aquabispora* may indicate treatment-specific microbial responses. Associations involving *Streptomyces*, *Epicoccum*, and Sordariales further suggest potential links to Fe nutrition and Fe-chelating activity. Together, these findings highlight complementary ecological patterns rather than contradictory results, although the functional roles of these taxa remain unverified. Interestingly, *V. paradoxus* inoculation did not significantly improve chlorophyll status or plant growth under Fe-deficient conditions in autoclaved soil (Supplementary Fig. S1), contrasting with the beneficial effects observed in nonsterile soil. This finding suggests that resident soil microorganisms may contribute to the growth-promoting effects of *V. paradoxus* under Fe limitation. However, soil autoclaving can also alter nutrient availability and soil physicochemical properties. Therefore, these results provide indirect evidence for a possible microbiome-mediated contribution rather than demonstrating a causal role of resident microorganisms in improved Fe nutrition.

### 4.4. Shared core microbiome and exploratory network analysis

Microbial co-occurrence networks can identify highly connected taxa that may be relevant to community organization (Agler et al., 2016). Our exploratory analysis identified *Funneliformis* and *Serendipita* among the highly connected fungal genera associated with *V. paradoxus* treatment under Fe limitation. Previous studies have reported beneficial roles of *Funneliformis* in nutrient acquisition and plant resilience (Shi et al., 2021), while members of *Serendipita* have been associated with enhanced nutrient acquisition, root development, and stress tolerance (Varma et al., 1999; Weiß et al., 2016). However, network connectivity represents statistical associations rather than direct microbial interactions or demonstrated functional contributions. Therefore, these taxa should be considered candidates for future functional validation. Core microbiome members are thought to provide functional resilience and maintain essential ecosystem processes despite environmental perturbations (Hanif et al., 2024; Toju et al., 2018). Core microbiome analysis identified genera consistently detected across treatments, including *Pseudomonas*, *Hydrogenophaga*, and *Funneliformis*. Their persistence indicates that these taxa remained prevalent despite treatment-associated differences in community composition. However, their presence alone does not establish functional resilience or contributions to plant Fe nutrition. *Pseudomonas* is a well-established plant-beneficial genus capable of siderophore production and induction of systemic resistance (Lugtenberg & Kamilova, 2009; Haas & Défago, 2005). Although evidence for direct plant growth promotion by *Hydrogenophaga* remains limited, it is frequently associated with healthy rhizospheres and nutrient cycling (Wang et al., 2025). However, *Funneliformis* is known for its role in nutrient exchange and microbiome restructuring (Shi et al., 2021). Thus, *V. paradoxus* inoculation appeared to modify selected community members while retaining potentially important resident taxa under Fe deficiency.

Treatment-related differences in microbial relative abundance may influence the inferred network structure. Thus, these genera should be regarded as candidates for independent validation rather than confirmed functional partners. Previous studies have reported beneficial roles of *Funneliformis* in nutrient acquisition and plant resilience (Shi et al., 2021), while members of *Serendipita* have been associated with enhanced nutrient acquisition, root development, and plant stress tolerance (Varma et al., 1999; Weiß et al., 2016). However, network connectivity represents statistical associations rather than direct microbial interactions or functional contributions. Therefore, these taxa should be considered putative beneficial fungal hubs whose roles in cotton Fe acquisition require experimental validation.

Indicator species analysis further identified *Shinella* and *Aquabispora* as markers of the community state associated with *V. paradoxus* treatment under indirect Fe deficiency. Indicator taxa respond consistently to specific environmental conditions and serve as markers of community assembly (De Cáceres & Legendre, 2009). Although the ecological role of *Aquabispora* in plant-associated environments remains poorly understood, its indicator status suggests a consistent association with the *V. paradoxus*-treated root community. Future studies should validate these microbial interactions under field conditions and determine the functional contributions of key helper and hub taxa. Exploratory Spearman correlations link selected taxa with Fe-related plant traits, although these associations do not demonstrate direct functional contributions. *Streptomyces* was positively associated with root Fe concentration and biomass, consistent with the documented capacity of members of this genus to produce siderophores and mobilize nutrients (Viaene et al., 2016). *Epicoccum* and members of Sordariales were associated with rhizosphere CAS-reactive Fe-chelating activity, suggesting potential links with Fe-related processes that require direct functional validation. Although these correlations do not establish causal relationships, they identify candidate taxa whose associations with Fe-related plant traits warrant validation in experiments with greater replication and direct functional testing.

## Conclusion

We found that *V. paradoxus* improves tissue Fe accumulation, photosynthetic performance, and plant growth in cotton exposed to bicarbonate-induced Fe limitation (Fig. 9). The lower root carbon concentration in *V. paradoxus*-inoculated plants was associated with inoculation, while bilateral inoculation produced greater improvements in growth and chlorophyll status than unilateral inoculation. However, the split-root experiment could not distinguish the effects of inoculum dose from root exposure or directly establish improved Fe acquisition. The absence of significant growth benefits in autoclaved soil further suggests that resident soil microorganisms may contribute to the beneficial effects of *V. paradoxus* under Fe deficiency. Although overall bacterial richness remained unchanged, *V. paradoxus* altered bacterial community composition and was associated with enrichment of selected taxa, including *Cellvibrio*, *Funneliformis*, and *Dominikia*. The persistence of shared core taxa such as *Pseudomonas*, *Hydrogenophaga*, and *Funneliformis*, together with exploratory identification of highly connected taxa associated with *V. paradoxus* treatment, suggests selective changes in community composition rather than broad shifts in microbial diversity. Exploratory associations of Streptomyces with root Fe accumulation and biomass, and of *Epicoccum* and Sordariales with CAS-reactive Fe-chelating activity, identify candidate taxa for future functional investigation. However, these associations do not establish a causal contribution of microbiome changes to Fe mobilization or uptake. Taken together, these findings highlight the potential of *V. paradoxus* to improve cotton Fe nutrition under bicarbonate-induced Fe limitation while identifying microbial taxa for future functional validation.

**Fig. 9.**
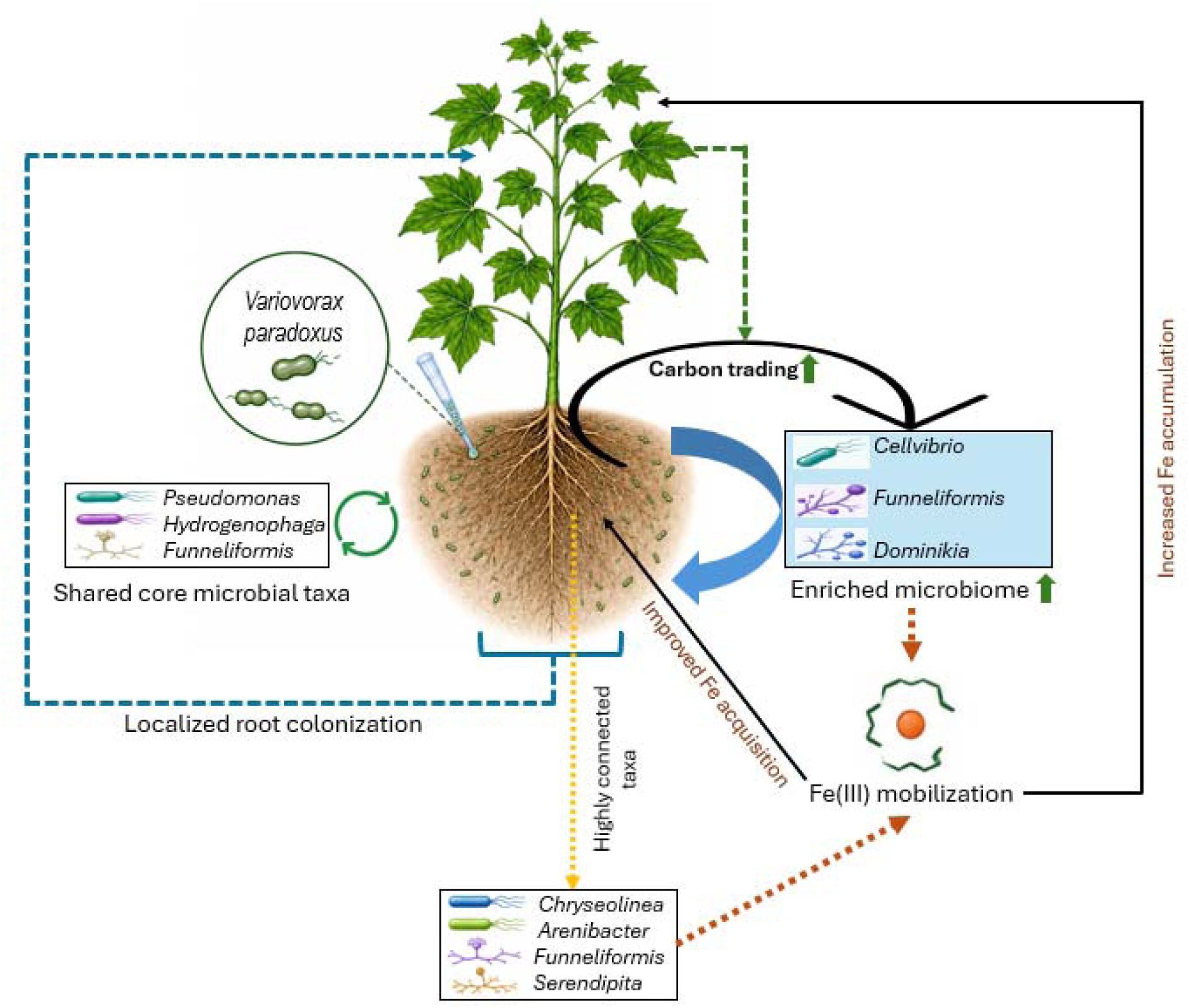
Conceptual model illustrating experimentally observed responses and hypothetical interactions associated with *V. paradoxus* inoculation under bicarbonate-induced Fe limitation in cotton. Observed responses include increased tissue Fe concentration, accompanied by change in root-associated microbial community composition and the relative abundance of selected microbial taxa. Potential contributions of these microbial changes to Fe acquisition remain hypothetical and require functional validation. Solid arrows represent experimentally supported responses, whereas dashed arrows indicate proposed relationships that were not directly demonstrated.

## Acknowledgements

The authors thank LC Sciences (Houston, TX, USA) for performing the 16S rRNA gene and ITS amplicon sequencing and for providing high-quality sequencing data.

## Data availability statement

The raw amplicon sequencing data generated in this study are publicly available in the NCBI BioProject database under accession numbers: 16S (PRJNA1497596) and ITS (PRJNA1497597).

