## Supplementary for "*Variovorax paradoxus* alleviates bicarbonate-induced Fe limitation in cotton by altering the root-associated microbiome"

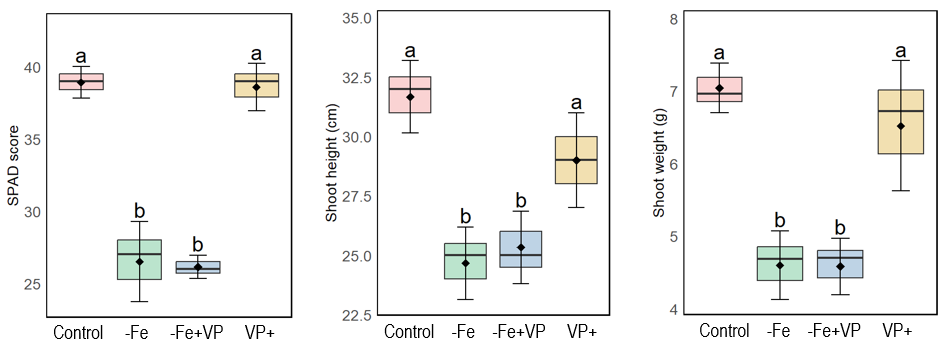

**Supplementary Fig. S1.** Effects of *V. paradoxus* (VP) inoculation on SPAD score, shoot height, and shoot weight of cotton plants grown in autoclaved soil under iron-deficient conditions for 6 weeks. Treatments included Control (Fe-sufficient), −Fe (iron-deficient), −Fe+VP (iron-deficient with VP inoculation), and VP+ (Fe-sufficient with VP inoculation). Boxplots represent three biological replicates (*n* = 3), with black diamonds indicating mean values. Different lowercase letters indicate significant differences among treatments according to Fisher’s LSD test (*p* < 0.05).
